# Encoding-linked pupil response is modulated by expected and unexpected novelty: Implications for memory formation and neurotransmission

**DOI:** 10.1101/2021.01.04.425232

**Authors:** Alex Kafkas

## Abstract

Whether a novel stimulus is expected or unexpected may have implications for the kind of ensuing encoding and the type of subsequent memory. Pupil response was used in the present study to explore the way expected and unexpected stimuli are encoded and whether encoding-linked pupil response is modulated by expectation. Participants first established a contingency relationship between a series of symbols and the type of stimulus (man-made or natural) that followed each one. At encoding, some of the target stimuli violated the previously established relationship (i.e., unexpected), while the majority conformed to this relationship (i.e., expected). Expectation at encoding had opposite effects on familiarity and recollection, the two types of memory that support recognition, and modulated differently the way pupil responses predicted subsequent memory. Encoding of unexpected novel stimuli was associated with increased pupil dilation as a predictor of subsequent memory type and strength. In contrast, encoding of expected novel stimuli was associated with decreased pupil response (constriction), which was predictive of subsequent memory type and strength. The findings support the close link between pupil response and memory formation, but critically indicate that this is modulated by the type of novelty as defined by expectation. These novel findings have important implications for the encoding mechanisms involved when different types of novelty are detected and is proposed to indicate the operation of different neurotransmitters in memory formation.

## 1. Introduction

Accurate prediction of the sequence of events may be seen as an attractive alternative to the uncertainty we so often face in everyday life. The human mind strives to form predictions about regularities in the environment (Friston et al., 2015), but this effort is disrupted when unexpected encounters occur that violate previous monotonies. However, expectedness, when new stimuli are detected and encoded into memory, can have consequences for the mechanisms that support learning and memory (Kafkas & Montaldi, 2015a, 2018a). Besides, expectation and especially unexpectedness is closely related to novelty (Kafkas & Montaldi, 2018b; Reichardt, Polner, & Simor, 2020) and the experience of surprise (Barto, Mirolli, & Baldassarre, 2013). Understanding the way expectation affects encoding of new stimuli is still lacking, but its exploration can benefit theories of learning and memory. Therefore, the interplay between expectation, novelty type and encoding of information is the focus of the present study.

As novelty can have diverse meanings (for reviews see Barto et al., 2013; Frank & Kafkas, 2020; Kafkas & Montaldi, 2018b; Ranganath & Rainer, 2003), in the present study, the term is used to refer to any experience or stimulus which lacks specific pre-existing representations from an episodic standpoint. In this sense, novelty may be based on features of the item itself (e.g., the picture of dice has not been presented before in the experiment, therefore the item is new) or on features of the context a stimulus is presented in (e.g., the picture of dice has not been presented, but its presentation is also surprising and unexpected in the current context). Finally, conceptual or semantic novelty, where a stimulus completely lacks pre-experimental conceptual representations as it happens, for example, in the case of abstract shapes or non-words, will not be examined here.

### 1.1. How does expectation define novelty and modulate memory encoding and retrieval?

Novelty detection itself is motivational (Bunzeck & Duzel, 2006) as a novel bit of information may indicate something significant in the environment that we may want to remember and integrate with existing knowledge, either to avoid it in the future or to reap possible rewards. However, the concept of novelty is not univocal in that different types of novelty can be discriminated each of which may have implications for the way we process, encode, act upon and remember them in the future (Frank & Kafkas, 2020; Kafkas & Montaldi, 2018b; Schomaker & Meeter, 2015). Novelty may enhance episodic memory formation directly as novel stimuli are prioritised at encoding (e.g., Geraci & Manzano, 2010; Tulving & Kroll, 1995) or indirectly as encoding benefits may ensue any stimulus presented in the context of novelty (e.g., Bunzeck & Duzel, 2006; Fenker et al., 2008; Kafkas & Montaldi, 2015a; Okuda, Højgaard, Privitera, Bayraktar, & Takeuchi, 2020; Schomaker, van Bronkhorst, & Meeter, 2014). However, the beneficial effect of novelty on memory is not absolute in that it does not always lead to subsequent memory boosts (e.g., Biel & Bunzeck, 2019). This indicates that novelty is prioritised in memory under certain conditions but not always.

Expectation may be an important dimension that allows different types of novelty to be discriminated. For example, an unexpected novel stimulus may trigger more rigorous encoding mechanisms (or attentional resource allocation) than an expected novel stimulus. However, the direction of episodic memory effects defined by the encoding of expected or unexpected stimuli is far from straightforward. For example, previous studies have revealed that stimuli conforming to previously established conceptual associations tend to be remembered better relative to incongruent stimuli (the congruency effect; e.g., Bartlett, 1932; Bein et al., 2015; Craik & Tulving, 1975; Staresina, Gray, & Davachi, 2008). In contrast, distinctive information that deviates from the perceptual or conceptual characteristics of the current context, and is therefore unexpected, has also been shown to result in enhanced memory encoding as revealed by greater subsequent memory retrieval (e.g., Rangel-Gomez & Meeter, 2013; Schmidt & Schmidt, 2017; Wallace, 1965). Furthermore, two recent studies (Greve, Cooper, Tibon, & Henson, 2019; Kafkas & Montaldi, 2018a), using expectation manipulations related to violation or validation of newly established rules, have found memory advantage for both expected and unexpected stimuli.

Greve et al., (2019) showed that a memory advantage can be seen not only for stimuli that are consistent with experimentally established arbitrary associations, but also for stimuli that violated this association (i.e., unexpected). Kafkas & Montaldi (2018a) showed that stimuli that conform to learned contingency rules (expected) and those that violate these rule (unexpected) are accompanied by increased memory performance based, nevertheless, on different types of memory. Expected stimuli were remembered better on the basis of familiarity, while unexpected stimuli were accompanied by increased recollection accuracy. Familiarity denotes memory for the item itself, without recall of associative details from the previous occurrence, whereas recollection indicates retrieval of associative information from the time of encoding (e.g., Kafkas & Migo, 2009; Kafkas & Montaldi, 2012; Mandler, 1980; Montaldi & Mayes, 2010; Yonelinas, 2002). Therefore, processing of expected and unexpected novel stimuli may implicate different encoding mechanisms, which they lead to qualitatively different memory experiences. The main aim in the present study was to explore whether encoding mechanisms triggered from the detection of expected and unexpected stimuli vary. To this end pupil response measures were employed during encoding and their relationship to subsequent type of memory were monitored adopting a subsequent memory effect protocol (Paller & Wagner, 2002).

### 1.2. Encoding-linked pupil responses and modulation by type of novelty

Before examining the specific hypotheses of this study, it is relevant to consider the evidence for the close link between episodic memory encoding and pupil response and what implications this link may have for encoding mechanisms. It has been shown that the pupil response at encoding can be predictive of the type and strength of ensuing memory (e.g., Kafkas & Montaldi, 2011, 2015a; Miller & Unsworth, 2019; Naber et al., 2013; Papesh, Goldinger, & Hout, 2012; Wetzel, Einhäuser, & Widmann, 2020). However, the studies appear inconsistent as to the pattern of pupil response that predicts subsequent memory. For example, increased pupil dilation for later remembered relative to forgotten stimuli has been reported (e.g., Miller & Unsworth, 2019; Papesh et al., 2012), but this effect appears to be more pronounced when contextual or unexpected novelty is detected (Kafkas & Montaldi, 2015a). On the other hand, other studies have shown that decreased pupil dilation or pupil constriction predict the type and/or the strength of subsequent memory (Kafkas & Montaldi, 2011; Naber et al., 2013; Wetzel et al., 2020). This finding is also consistent with the pupil old/new effect, whereby reduced levels of pupil dilation accompany new relative to old stimuli in recognition memory paradigms (e.g., Kafkas & Montaldi, 2012, 2015b; Otero, Weekes, & Hutton, 2011; Võ et al., 2008).

In an effort to account for the obvious inconsistency in the way pupil response reflects successful memory formation for novel stimuli, Kafkas and Montaldi (2018b), in a recent neurocognitive model, proposed that different neural pathways and neurotransmitter systems are tuned in to different types of novelty. Specifically, contextual, surprising and unexpected novelty modulates the noradrenergic sympathetic system resulting in dopamine and norepinephrine release in the hippocampus and increased pupil dilation. On the other hand, according to the same model, detection of absolute and expected novelty modulates the cholinergic parasympathetic system resulting in acetylcholine release in the hippocampus and pupil constriction patterns. Similar to the proposal by Kafkas & Montaldi (2018b), recent reviews (Joshi & Gold, 2020; Peinkhofer, Knudsen, Moretti, & Kondziella, 2019) have also highlighted the utility of pupil response patterns in understanding brain pathways involved in cognition.

### 1.3. The present study

In the present study, the encoding mechanisms engaged when expected and unexpected stimuli are encountered were explored and their effect on subsequent memory were measured. The pupil response was used as a measure of online processing during the encoding period. Overall, the experiment aimed at exploring how expectation modulates the encoding mechanisms engaged for successful (i.e., subsequent familiarity or recollection hits) and unsuccessful (i.e., a subsequently forgotten/missed items) memory formation. It is assumed that in the case of expected stimuli, only stimulus absolute novelty is in operation at encoding as the sequence of the event conform to the established rule, while a new stimulus is presented. On the other hand, in the case of unexpected stimuli, apart from stimulus absolute novelty, a degree of contextual novelty and surprise is in operation due to the violation of expected sequence of the event. Therefore, all stimuli used at encoding were experimentally novel (i.e., objects presented for the first time in the experiment), but a subset of them also violated contextual expectation.

The following specific hypotheses were explored: Based on findings from (Kafkas & Montaldi, 2018a), it was hypothesised that expected and unexpected stimuli will result in opposite effect on subsequent familiarity and recollection responses. Based on the memory-related pupil effects reported in previous studies (see 1.3.), it was hypothesised that pupil response patterns at encoding will be predictive of the type and possibly the strength of reported memory. However, consistent with the proposal from Kafkas and Montaldi (2018b), it was hypothesised that these pupil response patterns at encoding will be modulated by the expectation status of the stimuli resulting perhaps in different constriction or dilation patterns for expected and unexpected stimuli. Finally, whether the hypothesised pupil modulation by expectation only applies to subsequent recollections or it also affects subsequent familiarity responses, was also explored.

## 2. Material and Methods

### 2.1. Participants

A total of 40 participants gave informed consent and participated in the study, which received ethical approval from the University of Manchester Research Ethics Committee. From this original sample, 3 participants were excluded from all analyses due to incomplete/noisy eye tracking recordings (see 2.4 below; inclusion of their behavioural data did not change the reported memory effects). The final sample included 37 participants (34 females, 3 males) with a mean age of 19.32 years (SD = 2.75). The sample size was informed by previous studies with similar methodology (Kafkas & Montaldi, 2011, 2018a). Specifically, a power analysis using GPower tool (Faul, Erdfelder, Lang, & Buchner, 2007), determined that 36 participants is a sufficient sample to replicate the previously reported (Kafkas & Montaldi, 2018a) effect of expectation on memory (parameters: *η*^2^ = 0.11; effect size *f* = 0.35; power = 0.90). All participants had normal or corrected-to-normal (with contact lenses) vision and self-reported no history of psychiatric or neurological conditions. They were further asked to abstain from coffee and alcohol consumption 24 hours before participating in the study, while systematic use of psychotropic medicines and drugs were described as exclusion criteria.

### 2.2. Stimuli

A subset of stimuli from Kafkas and Montaldi (2018a) were used in the present study. Briefly, 246 simple man-made and natural items presented in grayscale were used in the different conditions of the experiment (size: 500 x 375 pixels; degrees of visual angle at presentation: 18.7h and 14.05v). As has been described in detail before (see Kafkas & Montaldi, 2011, 2015b), the stimuli were matched for luminosity and emitted luminance and were presented in grayscale, with low contrast on a uniform grey background (RGB = 130; Figure 1). These steps ensured that the stimuli are appropriate for pupillometry. The encoding and recognition tasks included 170 stimuli (85 man-made and 85 natural), while 72 stimuli (36 man-made and 36 natural) were used in the rule learning task (see 2.3.). Four stimuli were also used for practice. Finally, 6 symbols (line drawings) were used in the rule learning task and as contextual cues at encoding (see 2.3. and Figure 1).

**Figure 1.**
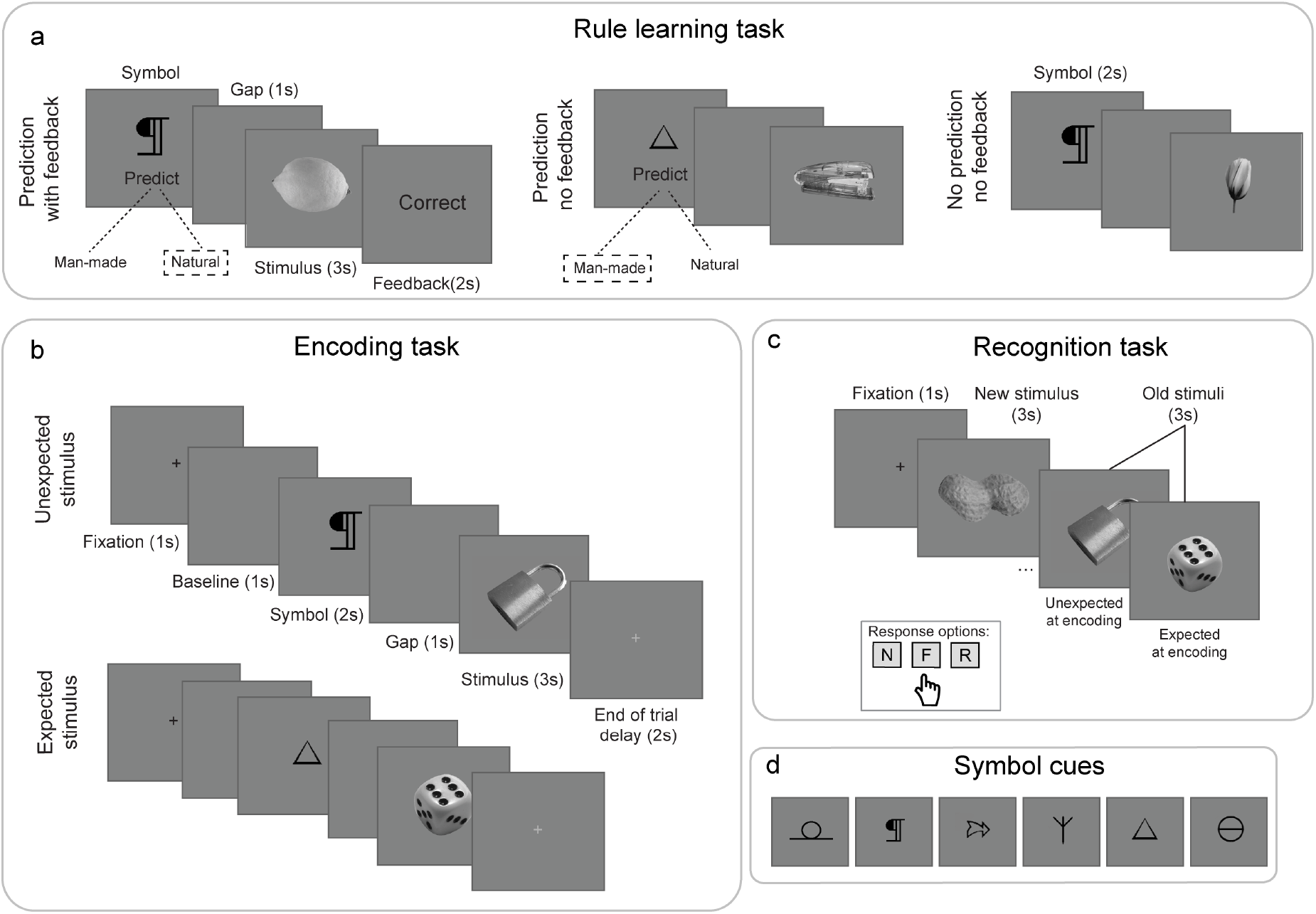
Experimental design. a) The rule learning task established contingency symbol-stimulus type associations, with each symbol (see d) consistently associated with either a man-made or a natural item (3 symbols assigned to each type). Twelve repetitions of the same symbol across the entire block were used. b) At encoding, a novel set of stimuli (completely different from those used in a) were presented after the same cuing symbols from the rule learning task. Sixty percent (60%) of the trials adhered to the established (in the rule learning task) rule (i.e., expected stimuli), while 40% of trials violated the previously learned rule (i.e., unexpected stimuli). The sequence within an expected and an unexpected event are illustrated in the examples. In the case of the unexpected stimulus (top sequence), the stimulus (padlock) is inconsistent with the rule established previously (i.e., symbol should be followed by a natural item). In the case of the expected stimulus (bottom sequence), the stimulus (dice) is consistent with the rule established previously (i.e., symbol should be followed by a man-made item). c) A recognition memory task tested memory for the previously encoded expected and unexpected stimuli intermixed with new (unstudied) foils. Participants pressed the corresponding button, while each stimulus was on the screen to report stimuli as either new (N), familiar (F) or recollected (R). d) All 6 symbols used as cuing stimuli in the experiment.

### 2.3. Procedure and design

Each experimental session consisted of 4 tasks: the rule learning, the encoding, the filled interval and the recognition task (Figure 1). The design of the experiment was similar to the design reported in Experiment 1a in Kafkas and Montaldi (2018a). The only difference was that eye tracking data were recorded at encoding and additional interstimulus interval was implemented to ensure recovery of pupil size from trial to trial (see below). The allocation of the object stimuli across the different conditions (rule learning, encoding, recognition) was completely random with the only restriction being the type of stimulus (man-made or natural) consistent with the expectation manipulation.

#### 2.3.1. The rule learning task

In the rule learning task participants learned a contingency relationship between a symbol and the type (man-made or natural) of the subsequent item. In each trial participants were asked to make a prediction as to whether a natural or a man-made object will follow a symbol (see Figure 1a). In total 6 symbols (Figure 1d) were used across this block and each of them was randomly assigned (across participants) to precede either a man-made (3 symbols) or a natural (3 symbols) item. The association remained the same throughout the rule learning task. In each trial the symbol remained on the screen until participants responded using two keyboard buttons to predict either a man-made or a natural item. After the participants’ prediction, a blank gray screen was presented for 1s followed by the object stimulus presented for 3s. Each symbol was repeated 12 times across the entire task, preceding a unique stimulus in each trial, which was nevertheless consistent with the specific symbol-stimulus type rule.

As shown in Figure 1a, for the first 36 trials (6 repetitions per symbol) a feedback screen followed the presentation of the stimulus, indicating whether the symbol-related prediction was correct or incorrect. For the next 18 trials no feedback was provided. Finally, the last 18 trials involved the same symbol-stimulus sequence, but participants were not asked to make a prediction. In these last trials the symbol remained on the screen for 2s. The sequence within the last 18 trials made the transition to the encoding task smoother. Overall, the design of this block ensured effective learning of the contingency relationships and the build-up of expectations based on the correct symbol-stimulus sequences (Kafkas & Montaldi, 2018a).

#### 2.3.2. The encoding task

Before participants started the encoding task, a standard nine-point calibration of the eye tracking system was performed, while each participant’s head was stabilised using an appropriate chinrest. Participants were informed that their eye movements would be recorded during this block to measure attention on the presented stimuli, but no explicit reference to pupil responses or to the type of memory test that would follow was made. This task was very similar to the final part of the rule learning task (last 18 trials; see 2.3.1) in that a symbol-stimulus sequence was adopted. The main difference was that in the encoding task participants were asked to focus on studying the stimuli carefully without the need to make any prediction in relation to the symbols. The same 6 symbols as in the rule learning task were used (20 trials each symbol), but a completely new set of man-made/natural stimuli (120 in total) were presented. The critical manipulation in this block was that only 60% of the trials (72 trials), spread across the entire encoding block, adhered to the previously established symbol-stimulus type relationship (expected stimuli). The remaining 40% of trials violated the symbol-stimulus rule and they were, therefore, unexpected. In the encoding task participants were asked to freely view the presented stimulus, without performing any other encoding task.

Each encoding trial started with a fixation cross (1s), followed by a blank gray screen, which served as a baseline period (1s) for calculating the trial-specific pupil response. The cuing symbol appeared for 2s followed by a blank screen (1s) and then the stimulus that remained on the screen for 3s. To ensure recovery of stimulus-related change in pupil size, a delay period of 2s was used after each stimulus and was marked with a gray fixation cross. Participants had the chance to take 2 self-paced breaks (every 40 trials) during this task.

#### 2.3.3. The recognition memory task

After completing the encoding task, each participant engaged in a filled interval of 10 minutes during which unrelated verbal and arithmetic tasks were performed. Subsequently, training and instructions pertaining the recognition memory decisions were offered. Occurrences of familiarity (F), recollection (R) and novelty (N) were explained and participants had the chance to practice these choices with 4 practice stimuli, two of which had been presented previously as practice items just before the encoding task. It was explained that a recollection response indicated the recall of additional contextual details associated with the encoding episode (e.g., a thought related to a specific stimulus). Contrary, a familiarity response meant that no additional detail came to mind related to the study episode apart from memory for the item itself. Similar to previous work with fMRI and eye tracking methodologies (e.g., Kafkas, Mayes, & Montaldi, 2020; Kafkas et al., 2017; Kafkas & Montaldi, 2012, 2014; Mayes et al., 2019) familiarity responses were provided using a 3-point rating scale from weak to strong familiarity. Participants explicitly instructed not to confuse confidence with the familiarity and recollection decision. Instead, it was stressed that they can be very confident they have encountered a stimulus before at encoding without being able to recollect additional details associated with this encounter (for previous work related to this important distinction see e.g., Kafkas et al., 2017; Kafkas & Montaldi, 2012; Mayes et al., 2019).

Before commencing the main part of this task, participants had the chance to ask questions in relation to the recognition decisions and they were also asked to explain to the experimenter what the response choices meant. This ensured appropriate use of the recognition memory decisions and accurate reporting of the experienced memory kind for each stimulus. Participants completed a total of 170 trials (120 from encoding and 50 new ones), each starting with a fixation cross. Each stimulus appeared for a maximum duration of 3s, during which participants were instructed to provide their recognition decision. This timing was also practiced in the practice trials before commencing the main block. No eye tracking data were recorded during this period and therefore a chinrest was not used.

#### 2.3.4. Debriefing

After the end of the experimental session participants were asked to explain what they thought the specific aim of the study was and were subsequently informed of the aims by the experimenter. As in the previous study in which a similar paradigm was used (Kafkas & Montaldi, 2018a), only a small number of participants (2) reported they realised the violation of the symbol-stimulus rule in some of the encoding trials. The two participants were included in all the analyses, but exclusion of their data did not change the direction of the main findings as reported here.

### 2.4. Apparatus, eye tracking recording and pre-processing

Apparatus: Each session was completed in a quiet testing room moderately lit (at about 250 lx) across all sessions and equipped with an ASL infrared eye tracking system (Applied Science Laboratories, Model Eye-Trac 6000; sampling rate 60 Hz). A standard-nine-point calibration was performed with each participant just before stating the encoding task. An artificial 4mm pupil was used at the end of encoding at the position of each participant’s eye to later convert the pupil size recordings into mm. All parts of the experiment were completed using a PC with a 19-inch monitor (resolution: 1280 x 1024). Participants had a distance of about 70cm from the computer monitor and a special chinrest was using for the period of eye tracking recording (encoding task). A standard computer keyboard was used to collect participants’ behavioural responses.

Pre-processing of the pupil data followed previously established protocols (e.g., Kafkas & Montaldi, 2011, 2015b). Specifically, blinks, artefacts and partial closures of the eyelid were identified on the pupil size recordings by the eye tracking software and were excluded from the analyses. Also, to ensure that no pupil trace was contaminated by other residual artefacts and noise, a grand mean was calculated across all pupil traces within each trial and pupil recordings diverging more than three standard deviations from the mean were also discarded. These traces are artefacts of the pupil recording mostly related to partial eyelid closures right before or after a blink. Trials with more than 50% of discarded traces were excluded from analyses and participants with too many excluded trials (more than 40% of total number) were removed from the sample (3 participants were removed; see 2.1.) As the discarded pupil recordings across all trials, in the final participant sample, were very low (less than about 3% of the total recorded traces), no interpolation was employed to replace these recordings. As in previous studies, (see e.g., Kafkas & Montaldi, 2015b), linear interpolation for the missing traces did not have any effect on the observed pupil effects and therefore analyses with non-interpolated data are reported below.

#### Pupil timeseries

The trial-related pupil recording lasted for 3s, which was the duration of each trial at encoding, giving therefore a total of 180 continuous pupil recordings per trial (if no blinks or artefacts were identified during this period). The average pupil size during a period of 1s preceding each symbol was used as trial-specific baseline. Pupil timeseries for each trial were calculated by subtracting the trial-specific baseline average from each one of the 180 pupil size recordings during the presentation of the target stimulus. Therefore, the pupil timeseries were expressed as deviations from a zero-point baseline allowing the discrimination of pupil dilation and constriction patterns. It is relevant to stress that the terms pupil dilation and constriction relate to variations in pupil size due to the presence of a stimulus relative to a preceding baseline period. They should not be considered as indicating an absolute match onto the physiological dilation and constriction responses of the pupil. That said, a reduction in pupil response is more likely to evince physiological constriction of the pupillary sphincter, while increased pupil response is more likely to relate to physiological dilation of the iris dilator muscle. Finally, for the analyses, the pupil timeseries were divided in 10 timebins of 300ms. These timeseries were the main pupil response measures reported in this experiment^1^.

### 2.5. Statistical analyses

Linear mixed effect analyses using maximum likelihood estimation were performed to analyse the effect of expectation and memory type on memory performance, response times at recognition (RTs) and pupil response at encoding. Memory performance was calculated using d’ (d-prime) separately for familiarity and recollection responses. These analyses were performed using the GAMLj module (Gallucci, 2019) in Jamovi 1.2.16 (The jamovi project, 2020), which implements linear mixed effect analysis based on the lme4 package (Bates, Mächler, Bolker, & Walker, 2015) in R software. An important advantage of modelling predictors (independent variables) and measures (dependent variables) using linear mixed effects, instead of analysing them using repeated measures ANOVAs, include the ability to control for the variance attributed to random factors, such as the participants in the sample or any other factor that may explain random variance across participants (Baayen, Davidson, & Bates, 2008). It is also better suited for unbalanced designs where the number of observations or available data may vary across participants or conditions (as in the present study).

In the linear mixed effects model for memory performance, expectation (expected versus unexpected) and subsequent memory type (familiar versus recollected) were set as fixed effect factors (predictors), while participants were specified as a random effect (random intercept). The same modelling approach was also adopted for the analysis of RTs. Similarly, to analyse the effect of expectation and memory type on encoding-related pupil response, expectation, memory type and time window at encoding were set as fixed factors, while the random factor of participants on the intercept was also fitted. The time variable included the pupil response over the period of encoding (3s) summarised into 10 timebins (moving window of 300ms). The memory type in this analysis included missed (M), familiar (F) and recollected (R) stimuli. For each analysis, type III Wald F tests (with Satterthwaite’s method for approximating degrees of freedom; Kuznetsova, Brockhoff, & Christensen, 2017; Satterthwaite, 1946) are reported and significant effects were further explored using pairwise comparisons with Bonferroni adjustment for multiple comparisons. For the main analyses, familiarity responses were collapsed across the three levels of rating, but they were also examined separately in the case of pupil response to explore the relationship between familiarity strength and pupil response.

To determine which random effects to include in the analyses, models with random effects of memory type and expectation status across participants (i.e., random slope of conditions across subjects) were fitted for comparison. These are summarised in Table 1. Including random slopes in mixed effects models has been shown to reduce type I and type II errors (Schielzeth & Forstmeier, 2009). The model that significantly improved the explained variance using fewer parameters included the memory type as random effect across participants for performance and RTs and both memory type and expectation as random effects for pupil response. The findings from these models are presented in the Results and the comparison parameters of the different hierarchical models from each analysis are summarised in Tables A.1-A.3 (Appendix A). For these comparisons, the likelihood ratio tests approach was adopted, using the *χ*^2^ distribution, which compares whether the addition of a parameter (in this case a random effect) significantly improves the fit of the model. Additionally, changes in the variance explained by the models when fixed and random effects are included were also taken into account using a pseudo R^2^ measure (Johnson, 2014; Nakagawa & Schielzeth, 2013). Specifically, R^2^_marginal_ indicates the variance explained by the fixed effects only relative to the total variance of the dependent measure, while R^2^_conditional_ indicates the variance explained by the fixed and random effects together relative to the variance in the dependent measure. These metrics are also provided in the description of the results as indirect indicators of effect size. A significance level of *p* < 0.05 was adopted for all analyses.

**Table 1.**
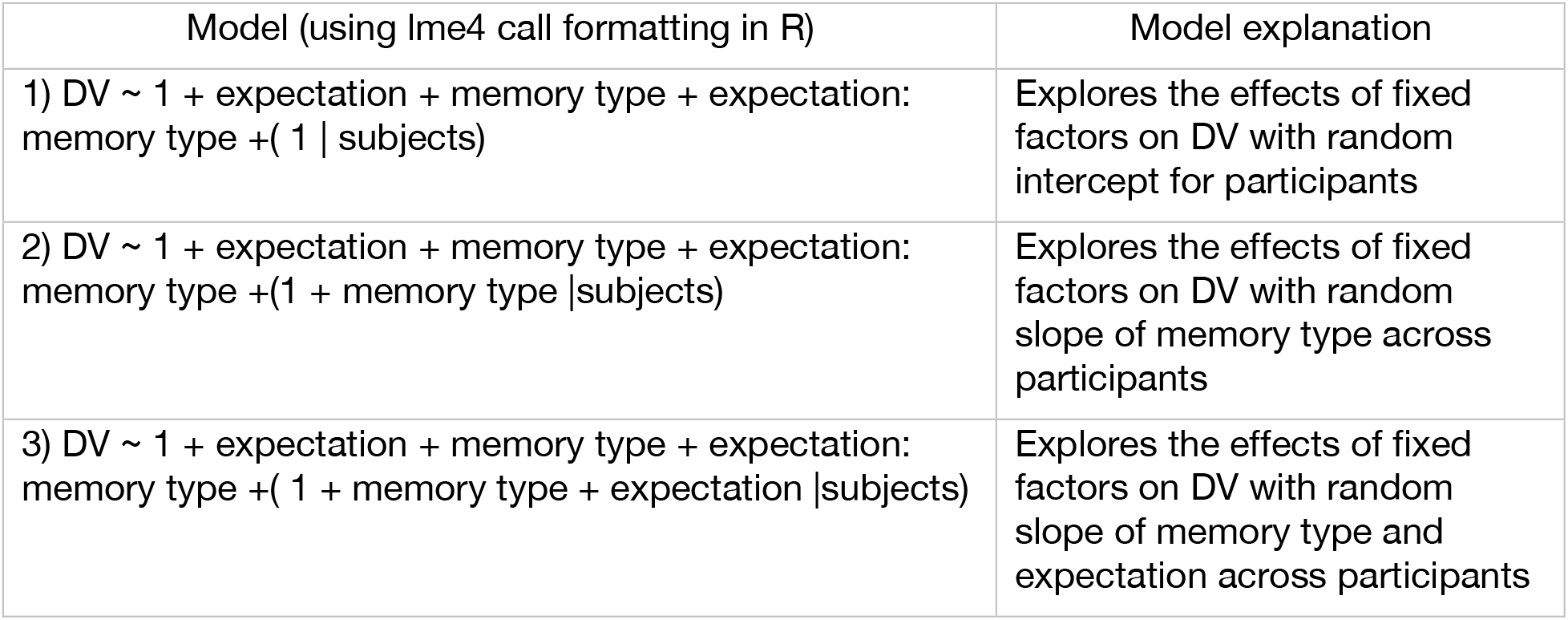
Three nested models with the same fixed factors (expectation, memory type and expectation by memory type interaction) but increased complexity in the included random factor for the analysis of the dependent variables (DV)

## 3. Results

### 3.1. Behavioural effects

#### 3.1.1. Rule learning

Mean accuracy across participants and symbol repetition in the rule learning task was 0.86 (SD = 0.12). Importantly, as shown in Figure 2, symbol prediction accuracy significantly increased with repeated presentations of the symbols (*F*_11,396_ = 27.6, *p* < 0.001), while RTs were significantly reduced across symbols repetitions (*F*_11,396_ = 7.86, *p* < 0.001). Therefore, the rule learning task was successful at creating clear expectations regarding the symbol-stimulus sequences.

**Figure 2.**
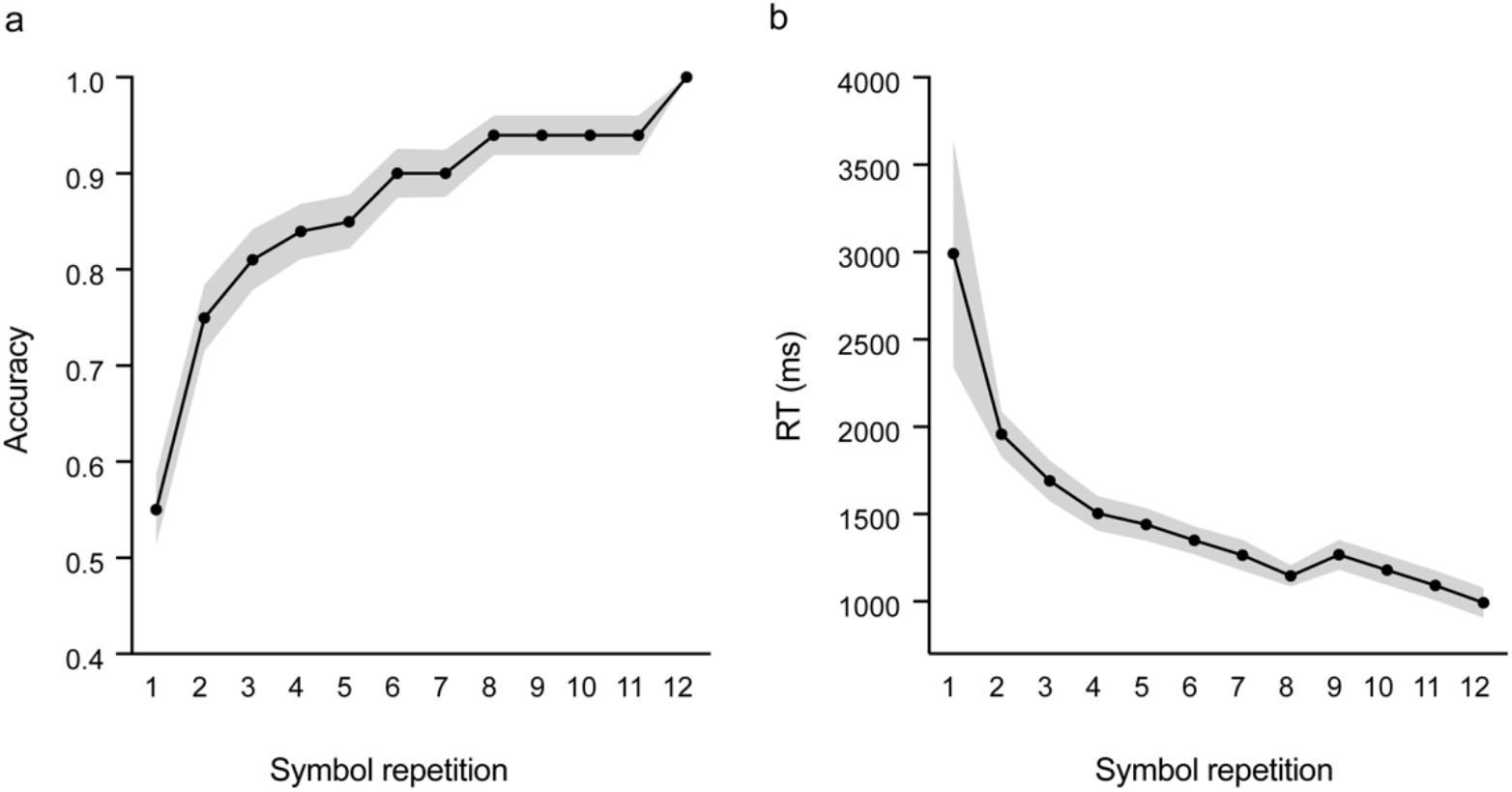
Prediction accuracy and response times (RTs) across 12 repetitions of each symbol in the rule learning task. Significant improvement in accuracy and reduction in RTs indicate successful learning of the contingency associations and therefore development of clear expectations for symbol-stimulus sequences.

#### 3.1.1 Differential effect of expectation on memory performance for familiarity and recollection

A linear mixed effect analysis was conducted to explore the effect of expectation status at encoding (expected versus unexpected) and type of subsequent memory (familiar versus recollected) on memory performance (measured using d΄). As can be seen in Table A.1, the model that included the random effect of participants (intercept) and the random effect of memory type significantly improved the data fit over the simpler (fixed effects + random effect of participants) or the more complex model (fixed effects + random effects of participants, memory type and expectation). This model explained large proportion of variance in memory performance (R^2^_marginal_ = 0.30; R^2^_conditional_ = 0.77) and significantly improved the fit. This means that the magnitude of the effect of memory type on memory performance varies across participants and therefore their random slopes should be modelled in the analysis of fixed effects presented below.

The mean proportions across the different response outcomes (hits, false alarms, misses and correct rejections) for expected and unexpected stimuli at encoding are summarised in Table 2. The analysis showed that the expectation status of stimuli at encoding did not predict overall subsequent memory performance (*F*_1,74_ = 0.91, *p* = 0.34), however, significantly higher performance associated with familiarity responses relative to recollection responses (*F*_1,37_ = 24.62, *p* < 0.001, *β* = −0.75, SE = 0.12). Importantly, the expectation by memory type interaction was significant (*F*_1,74_ = 44.49, *p* < 0.001) indicating that differences in memory performance for familiarity and recollection responses varied across expectation status at encoding (*β* = 0.83, SE = 0.12). Specifically, post hoc tests showed that familiar responses to expected stimuli were associated with increased memory performance relative to familiar responses to unexpected stimuli (*t*_76.1_ = 5.32, *p*_*bonferroni*_ < 0.001). Opposite to this effect, recollection responses to unexpected stimuli were associated with increased memory performance relative to recollections for expected stimuli (*t*_76.1_ = −3.99, *p*_*bonferroni*_ < 0.001; Figure 3). Therefore, the findings indicate that the expectation status at encoding has opposite effects on memory performance for familiarity and recollection responses, with the latter favouring unexpected stimuli and the former favouring expected stimuli. Similar effects were found when d’ was calculated using the criterion of independence for familiarity responses (Yonelinas & Jacoby, 1995) or when using the proportion of hits minus the proportion of false alarms as a measure of memory performance.

**Table 2.** Mean proportions and RTs for the different response outcomes at recognition for expected and unexpected (at encoding) stimuli Note: Numbers in the parentheses are standard deviations. F_hits_=familiarity hits; R_hits_ = recollection hits; M = misses; FA = false alarms; F_FA_ = familiarity false alarms; R_FA_= recollection false alarms; CR= correct rejections. R_hits_ are means across 22 participants who reported recollections, while the rest of the proportions are means across 37 participants^2^.

| Response | Proportions |  | RTs (in ms) |  |
| --- | --- | --- | --- | --- |
|  | Expected | Unexpected | Expected | Unexpected |
| $F_{\text{hits}}$ | 0.82(0.09) | 0.67(0.13) | 1354(201) | 1362 (210) |
| $R_{\text{hits}}$ | 0.10(0.05) | 0.20(0.06) | 1359(300) | 1341 (296) |
| M | 0.12(0.08) | 0.20(0.10) | 1427(316) | 1335 (329) |
| $F_{\text{FA}}$ | 0.28(0.16) | | 1404(281) | 1401 (271) |
| $R_{\text{FA}}$ | 0.01(0.03) | | 1425(368) | 1379 (508) |
| CR | 0.72(0.16) |  | 1272(217) | 1272 (217) |

Note: Numbers in the parentheses are standard deviations. $F_{\text{hits}}$ =familiarity hits; $R_{\text{hits}}$ = recollection hits; M = misses; FA = false alarms; $F_{\text{FA}}$ = familiarity false alarms; $R_{\text{FA}}$ = recollection false alarms; CR= correct rejections. $R_{\text{hits}}$ are means across 22 participants who reported recollections, while the rest of the proportions are means across 37 participants<sup>2</sup>.

**Figure 3.**
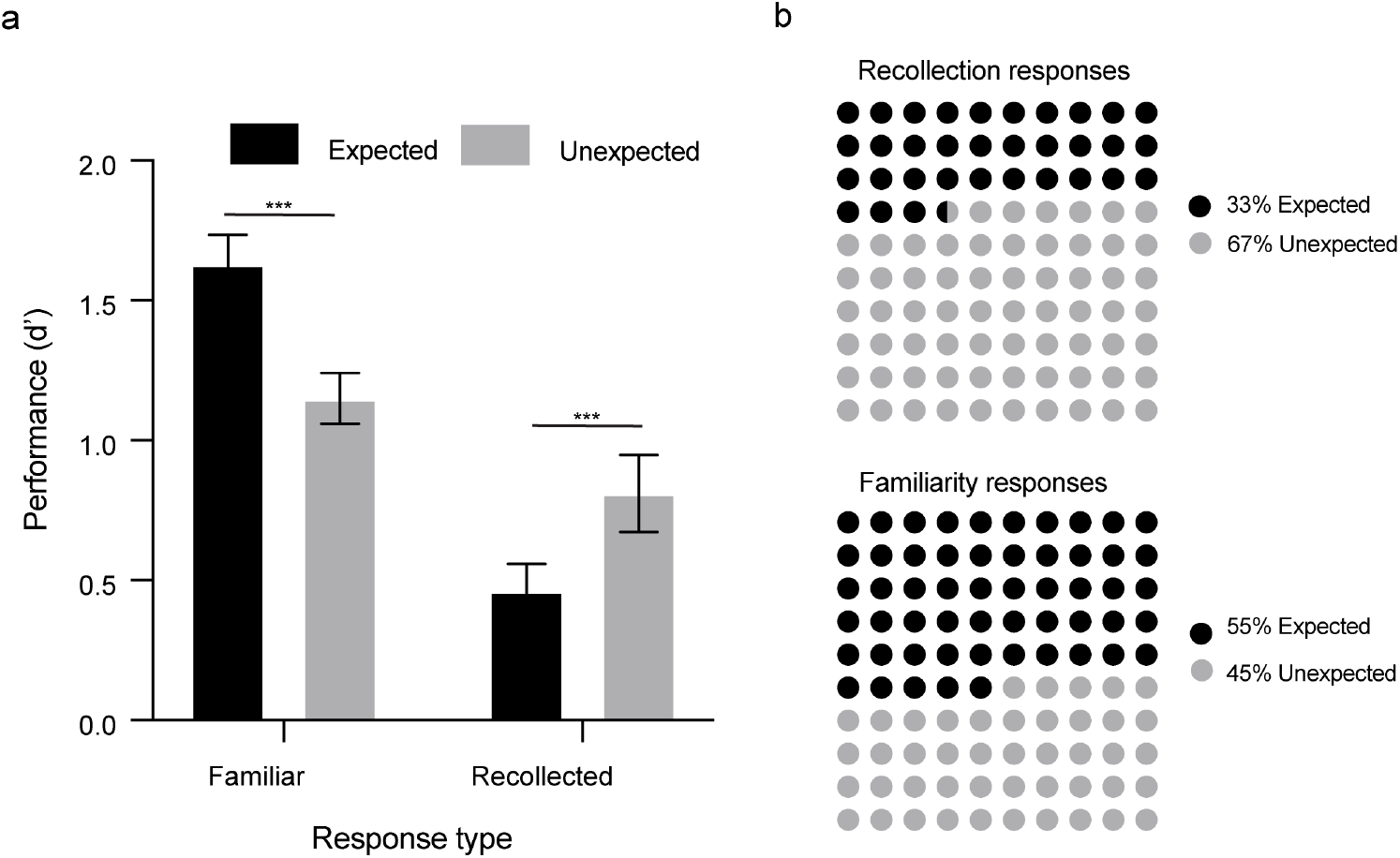
Novel expected and unexpected stimuli at encoding have opposite effects on familiarity and recollection responses at retrieval. a) Memory performance (d’) for subsequent familiarity and recollection responses for stimuli encoded as expected or unexpected. Error bars show standard error of the mean. b) Distribution of subsequent familiarity and recollection responses for expected and unexpected stimuli at encoding. Percentages represent division of the total number of recollection (upper panel) and total number of familiarity responses (lower panel) across expected and unexpected stimuli.

#### 3.1.2. RTs analysis

RTs were analysed in the same way as memory performance. Inclusion of memory type as a random effect significantly improved the model fit (for the different metrics and comparisons across the 3 models; see Table A.2), resulting in increased R^2^_conditional_ of 0.84. The R^2^_marginal_ of 0.008 shows that the fixed factors do not reliably predict the variance in RTs. Indeed, neither the main effect of expectation (*F*_1,53.3_ < 1) nor the main effect of response (*F*_1,23.6_ = 1.00, *p* = 0.32) nor the expectation by response interaction (*F*_1,53.9_ = 1.24, *p* = 0.27) were significant. Therefore, the expectation status at encoding and the type of memory at recognition did not have any effect on the RTs at recognition.

### 3.2. Pupil response at encoding by stimulus expectation and subsequent memory type

A linear mixed effect analysis was conducted to explore the effect of expectation status at encoding (expected versus unexpected), memory type (M, F, R) and time (encoding duration divided in 10 time-bins of 300ms) on pupil response. As can be presented in Table A.3, the model that included the random effect of participants (intercept), memory type and expectation (i.e., random slope of conditions of interest on pupil response) significantly improved the data fit (R^2^_marginal_ = 0.24; R^2^_conditional_ = 0.46). This means that the magnitude of the effect of memory type and expectation on pupil response varies across participants and therefore their random slopes should be modelled in the analysis of fixed effects presented below.

The analysis showed that the pupil response at encoding was significantly higher for unexpected relative to expected stimuli (*F*_1,39.8_ = 52.35, *p* < 0.001; *β* = 0.06, SE = 0.008), but it did not differ significantly across the three memory types (*F*_2,29_ = 2.03, *p* = 0.150). However, the memory type at retrieval significantly predicted the degree of pupil response at encoding, but the shape of this effect was different depending on the expectation status of stimuli at encoding. This finding was qualified by the significant expectation by memory type interaction (*F*_2,1791_ = 121.54, *p* < 0.001) and the significant triple interaction between expectation, memory type and time (*F*_18,1742_ = 11.18, *p* < 0.001). As shown in Figure 4, for expected stimuli there was a significant linear reduction of pupil response during encoding as a function of subsequent memory type (across M, F, R; *β*_*linear*_ = −0.05, SE = 0.009, *p* < 0.001). In contrast, for the unexpected stimuli, a significant linear increase of pupil response during encoding was associated with later memory type (across M, F, R; *β*_*linear*_ = 0.08, SE = 0.009, *p* < 0.001). This differential linear trend of the pupil response by the subsequent memory type, depending on the expectation status of stimuli, became significant half-way through the encoding period around 1500ms after stimulus onset for both expected (*β* = −0.04, SE = 0.081, *p* = 0.03) and unexpected stimuli (*β* = 0.04, SE = 0.019, *p* = 0.02), and remained significant until the end of the encoding period (i.e., 3s). Similar differential linear effects were also found when examining subsequent memory type separated by familiarity strength (i.e., M, F1, F2, F3, R) for expected (*β* = −0.05, SE = 0.009, *p* < 0.01) and unexpected stimuli (*β* = 0.08, SE = 0.009, *p* < 0.01). The only difference in the effect was that the linear increase in pupil response across memory type became significant slightly later (1800ms) in the case of unexpected stimuli.

**Figure 4.**
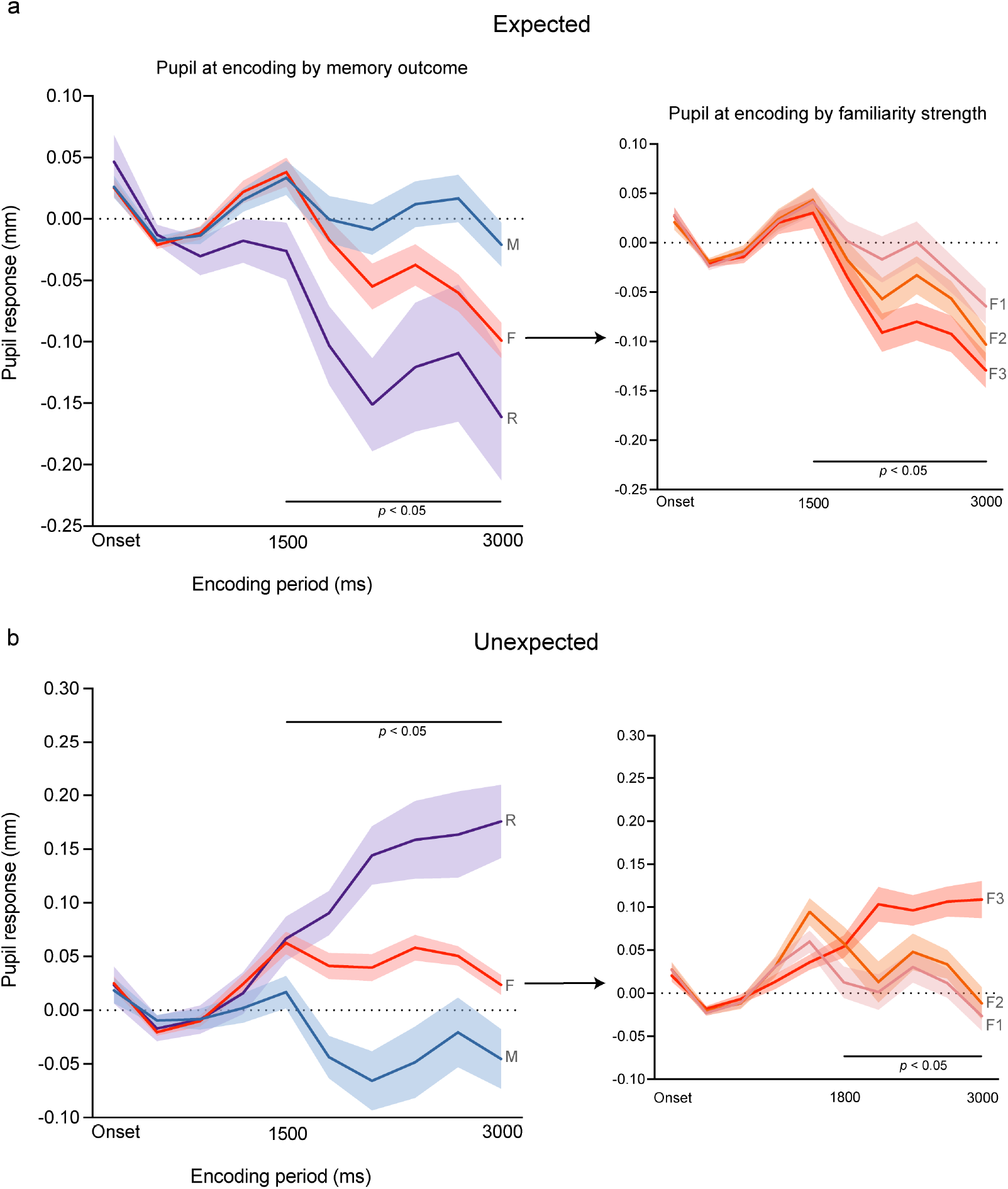
Pupil response at encoding across expected (a) and unexpected stimuli (b) as a function of subsequent memory type and familiarity strength. M = missed old stimuli; F = familiarity hits collapsed and separately across F1 (weak), F2 (moderate) and F3 (strong) familiarity; R = recollected hits. The degree of pupil response at encoding significantly predicted subsequent type of memory (or memory strength), but with opposite directions according to the expectation status of the encoded stimuli. The period during which the linear effect of subsequent memory on pupil response is significant (*p* < 0.05) is marked.

## 4. Discussion

In the present study, pupil response was examined during the encoding of expected and unexpected novel stimuli. The results replicated the previously reported finding (Kafkas & Montaldi, 2018a) that the encounter of expected and unexpected stimuli have contrasting effect on familiarity and recollection, the two types of memory that support recognition memory. Importantly, it was shown that the pupil response patterns at encoding are modulated by the expectedness of the encoded stimuli. Specifically, expected stimuli that conform to previously established regularities, were accompanied by a linear reduction (or constriction) of pupil response tracking the type and the strength of subsequent memory. On the other hand, encoding of unexpected stimuli that disrupt contextual or temporal regularities in the sequence of events, resulted in increased pupil response patterns (or dilations) predicting the strength and the type of subsequent memory.

This mirror-image pattern following the detection of expected and unexpected novel stimuli is consistent with the hypothesis that distinct encoding mechanisms are involved when the expectedness of novelty varies (for this proposal see Kafkas & Montaldi, 2018b). These expectation-modulated encoding mechanisms, as revealed by the pupil response patterns, were not only relevant for recollection, but also affected memory formation resulting in familiarity. The implications of these findings for novelty encoding, memory performance and the utility of pupil response as indicator of memory formation are discussed below. Also, in line with proposals that pupil response patterns provide a window into the involved neural circuitry (Joshi & Gold, 2020; Kafkas & Montaldi, 2018b) and the known relationship between sympathetic and parasympathetic pathways in controlling dilation and constriction patterns (Steinhauer & Hakerem, 1992; Steinhauer, Siegle, Condray, & Pless, 2004; see also Widmann, Schröger, & Wetzel, 2018), the involvement of different neurotransmitter systems in encoding expected and unexpected novelty are also discussed.

### 4.1. Expectation, memory formation and implications for type of memory

The behavioural findings show that whether a stimulus is expected or unexpected has consequences for the type of memory that is formed, which is reflected subsequently when memory is probed. Expected stimuli, that conform to established rules, are more likely to be remembered on the basis of familiarity, whereas unexpected stimuli, that violate the established rule, are more likely to produce accurate recollections. This opposite effect of expectation on the kind of memory supporting recognition is comparable to the previously reported effect using a similar paradigm (Kafkas & Montaldi, 2018a). However, in the previous paradigm, apart from the expectation status of stimuli, the type of encoding also varied. In the present study, the effect of expectation was replicated without requiring different encoding tasks, confirming, therefore, that the differential effect of expectation on later memory type cannot be explained by decision variations at encoding.

To the extent that unexpected stimuli are more distinctive at encoding, we may assume that these would receive preferential attentional allocation (Schmidt & Schmidt, 2017). This suggestion agrees with the general pattern of pupil response that accompanied unexpected stimuli at encoding. Traditionally, increased pupil dilations are thought to indicate enhanced resource allocation, enhanced processing and effort (e.g., Granholm, Asarnow, Sarkin, & Dykes, 1996; Hess & Polt, 1964; Kahneman, 1973; see discussion of pupil patterns below in 4.2.). But is this selective attentional allocation to the distinctive unexpected items at the expense of expected stimuli in terms of subsequent memory performance? Based on the present findings, the answer has to be twofold as it depends on the type of memory that is formed and supports retrieval. When examining recollection alone, there is a clear advantage for unexpected stimuli, therefore a presumed selective attentional allocation at encoding appears to be advantageous for the distinctive (unexpected) items. Consistent with this, recent neurocomputational evidence (Frank, Montemurro, & Montaldi, 2020) suggests that detection of unexpected stimuli promotes pattern separation in the hippocampus, a computational mechanism that supports subsequent recollection (Azab, Stark, & Stark, 2014; Bakker, Kirwan, Miller, & Stark, 2008). Nevertheless, selective allocation of attention to the unexpected stimuli does not benefit familiarity discrimination of these stimuli. Instead the expected stimuli are the ones benefited by a boost in familiarity memory.

These findings may have important educational implications, in that learning objectives should optimally combine scenarios in which both expected and unexpected new bits of information are offered to learners. Therefore, a learning session may have optimal outcomes, in terms of memory retention, when information conforms to previously established rules. On the other hand, unexpected and surprising information, for example discussing a concept from a novel, unexpected or less obvious standpoint, can be beneficial for the creation of elaborative memories. These memories can be optimally benefitted from the creation of new associative links.

Taken together, the findings indicate that inconsistences in the way expectation affects memory can be resolved by examining the type of memory that is formed and retrieved. Indeed, this tug of war between expected and unexpected stimuli is resolved by involving different encoding mechanisms. As discussed in the next section, this is closely demonstrated by differential pupil response patterns.

### 4.2. Type of novelty modulates pupil response patterns at encoding

The pupil response at encoding was closely related to subsequent memory type differentiating between missed, familiar and recollected stimuli. It also differentiated between multiple levels of memory strength, when the full scale of familiarity strength was used. However, the way the pupil response relates to subsequent memory was not unidirectional, but the pattern was modulated by expectation. Previous studies have provided inconsistent findings with respect to the direction of encoding-linked pupil responses, with some studies proposing increased dilation patterns as predicting successful memory formation (e.g., Papesh et al., 2012), while others finding that the degree of diminished dilation or constriction predicts successful memory formation (e.g., Kafkas & Montaldi, 2011; Naber et al., 2013).

The present findings unite these observations as they clearly suggest that the pattern of pupil response at encoding relates to the detection of different types of novelty. Therefore, unexpected stimuli are more likely to result in pupil dilation patterns and, importantly, the degree of this dilation predicts the strength and type of subsequent memory. In contrast, expected stimuli result in greater pupil constriction patterns at encoding and, critically, the degree of this constriction relates to the strength and type of reported memory. An interesting observation is that differential memory advantage between expected and unexpected stimuli can occur in quick succession during memory formation within the same experimental session. Therefore, the encoding mechanisms that prioritise different types of novelty, rely on fast switching to accomplish flexible memory formation.

A free-viewing task was used at encoding as it is considered more appropriate for pupillometry studies exploring memory formation (see also extensive discussion in Kafkas & Montaldi, 2011). Considering the sensitivity of the pupil response to effortful processing and working memory load (Kahneman & Beatty, 1966), it is undoubtable that when a decisional task is employed, an effect on pupil response may be found, not necessarily because a memory for the stimulus is formed, but because a decision is being made. A recent study proposed that pupil response patterns at encoding (and especially pupil dilations) are not predictive of subsequent memory but only reflect decision urgency (Gross & Dobbins, 2020). The findings here show that this explanation is limited and certainly does not apply to all reported encoding-linked pupil effects. Effort associated with a cognitive task at encoding or the urgency of taking a decision, may infiltrate the pupil response, but they cannot be the sole explanation of the close link between pupil response and successful memory formation. Instead, the findings show that the most critical factor explaining previous inconsistencies in the way pupil response relates to subsequent memory is the type of novelty detected.

### 4.3. Implications for novelty mechanisms, brain networks and neurotransmission

What is the cognitive origin of these effects and what predictions can be made regarding their neural underpinnings? Expected stimuli are more likely to be encoded in a way that supports familiarity with the item, benefited by the detection of highly predictive outcomes. Consistent with the visual discrimination literature, where predictive cues result in better visual discrimination of stimuli (e.g., Bollinger, Rubens, Zanto, & Gazzaley, 2010; Posner, Snyder, & Davidson, 1980; Puri & Wojciulik, 2008; Trapp, Lepsien, Kotz, & Bar, 2016) expected items at encoding drive more fluent and efficient processing of information. Nevertheless, the degree of novelty, even in the case of expected outcomes, is sufficient to involve encoding mechanisms in the MTL (Kafkas & Montaldi, 2014). At the same time, this type of novelty detection presumably results in increased release of acetylcholine via the implication of the parasympathetic system (which is exclusively cholinergic) resulting in the observed pupil constriction patterns (Loewenfeld, 1999; Steinhauer et al., 2004) (for the role of acetylcholine in learning, memory and plasticity see Haam & Yakel, 2017; Hasselmo, 2006). Therefore, it is proposed here that the detection of expected novel stimuli enables encoding via cholinergic input to the MTL (and especially the cortical regions along the parahippocampal gyrus; see Kafkas et al., 2017) and makes them more easily recognisable on the basis of familiarity when encountered again.

On the other hand, unexpected stimuli are more surprising within a given context leading to aroused attention and enhanced exploratory behaviour (Barto et al., 2013). For example, unexpected stimuli are more likely to result in increased visual fixations (Kafkas & Montaldi, 2015a), and, like in the presented study, increased levels of pupil dilation (Kafkas & Montaldi, 2015a). At the neural level, contextual/unexpected novelty produces enhanced P300 event‐related brain potential (ERP; Polich, 2007; Ranganath & Rainer, 2003), which also relates to pupil dilation (Murphy, Robertson, Balsters, & O’connell, 2011) and has been linked to hippocampal function (Fonken, Kam, & Knight, 2020). Importantly, it has been shown to engage a functional loop involving the dopaminergic midbrain (especially the SN/VTA) and the hippocampus (Kafkas & Montaldi, 2015a; Lisman & Grace, 2005; Shohamy & Wagner, 2008; Wittmann, Bunzeck, Dolan, & Duzel, 2007). The adjacent locus coeruleus (LC), which is associated with the degree of phasic pupil dilation (Eldar, Cohen, & Niv, 2013; Mather, Clewett, Sakaki, & Harley, 2016; Murphy, O’Connell, O’Sullivan, Robertson, & Balsters, 2014) has been linked to the detection of saliency, novelty and the attentional prioritisation of significance in the environment (Aston-Jones & Cohen, 2005; Markovic, Anderson, & Todd, 2014; Mather et al., 2016; Vazey, Moorman, & Aston-Jones, 2018). It is also a hotspot of norepinephrine (Szabadi, 2013; Walling, Brown, Milway, Earle, & Harley, 2011), as well as dopaminergic (Duszkiewicz, McNamara, Takeuchi, & Genzel, 2019; Kempadoo, Mosharov, Choi, Sulzer, & Kandel, 2016; Takeuchi et al., 2016; Wagatsuma et al., 2018) release to the hippocampus and the rest of the MTL. Therefore, it is proposed here that the pupil dilation as predictor of memory formation is linked to the detection of unexpected novelty and denotes dopaminergic and/or noradrenergic modulation (from SN/VTA and LC) of the hippocampus and the MTL cortex. This mechanism makes it more likely that unexpected stimuli are initially pattern separated and subsequently recollected.

Although in the present study expectation is proposed as a central factor defining the encoding mechanisms engaged when novel (unstudied) stimuli are presented, the proposal does not preclude effects of expectancy on processing non-novel stimuli (e.g., recently pre-familiarised items; for relevant evidence see Frank, Kafkas, & Montaldi, 2020; Kafkas & Montaldi, 2015a, 2018a). Therefore, the present findings and previous observations suggest that expectancy has a broad role in guiding memory encoding, reencoding as well as memory search and retrieval.

### 4.4. Conclusions

Taken together, the present findings demonstrate that the close link between pupil response and memory formation is sensitive to the type of novelty as defined by expectation. It is proposed that the relationship between either pupil dilation or constriction patterns, on the one hand, and the type of subsequent memory on the other, denotes the employment of different encoding mechanisms. These mechanisms are proposed to exert differential neurotransmitter control on memory formation regions of the brain and occur in quick succession in the service of flexible memory formation.

## Footnotes

1 Average or peak pupil responses can be used to summarise the trial-related pupil changes but their utility is limited as they do not allow evaluation of the time course of pupil change during the information processing period.

2 This is the reason the proportions of old stimuli (F_hits_ + R_hits_ + M) do not add-up to 1.00 as R_hits_ are calculated for N = 22 (participants reporting Rs), while F_hits_ and M are calculated for N = 37.

## Notes

### Competing Interest Statement

The authors have declared no competing interest.

