## Supplementary material for "Encoding-linked pupil response is modulated by expected and unexpected novelty: Implications for memory formation and neurotransmission": Table A.

**Alex Kafkas**
School of Biological Sciences, Division of Neuroscience & Experimental Psychology, University of Manchester, UK

**Table A.1.** Model comparisons across three nested models with the same fixed factors (expectation, memory type and expectation by memory type interaction) but increased complexity in the included mixed effects for the analysis of memory performance (d').

| Model (using *lme4* call formatting in *R*) | $R^2_{marginal}$ | $R^2_{conditional}$ | AIC | ICC | Random Effect LRT |
|---|---|---|---|---|---|
| 1) d ~ 1 + expectation + memory type + expectation: memory type +( 1 \| subjects) | 0.30 | 0.39 | 304.42 | 0.12 | $\chi^2(1) = 2.84$, $p = 0.09$ |
| 2) d ~ 1 + expectation + memory type + expectation: memory type +(1 + memory type \| subjects) | | 0.77 | 265.72* | 0.43 | $\chi^2(2) = 42.7$, $p < 0.001$ |
| 3) d ~ 1 + expectation + memory type + expectation: memory type +( 1 + memory type + expectation \| subjects) | | 0.78 | 269.03 | 0.45 | $\chi^2(3) = 2.69$, $p = 0.44$ |

Note: $R^2_{marginal}$ is the same across all models as the fixed effects remain the same; AIC = Akaike information criterion (smaller numbers indicate preferred model); ICC = intraclass correlation coefficient; *best model fit based on variance explained, significance of improvement after including the random effect and the parsimony of the model indicated by AIC.

**Table A.2.** Model comparisons across three nested models with the same fixed factors (expectation, memory type and expectation by memory type interaction) but increased complexity in the included mixed effects for the analysis of response times.

| Model (using *lme4* call formatting in *R*) | $R^2_{marginal}$ | $R^2_{conditional}$ | AIC | ICC | Random Effect LRT |
|---|---|---|---|---|---|
| 1) RT ~ 1 + expectation + memory type + expectation: memory type +( 1 \| subjects) | 0.008 | 0.59 | 1512.40 | 0.58 | $\chi^2(1) = 41.7$, $p < 0.001$ |
| 2) RT ~ 1 + expectation + memory type + expectation: memory type +(1 + memory type \| subjects) | | 0.84 | 1486.50* | 0.80 | $\chi^2(2) = 29.9$, $p < 0.001$ |
| 3) RT ~ 1 + expectation + memory type + expectation: memory type +( 1 + memory type + expectation \| subjects) | | 0.84 | 1492.28 | 0.80 | $\chi^2(3) = 0.22$, $p = 0.97$ |

Note: $R^2_{marginal}$ is the same across all models as the fixed effects remain the same; AIC = Akaike information criterion (smaller numbers indicate preferred model); ICC = intraclass correlation coefficient; *best model fit based on variance explained, significance of improvement after including the random effect and the parsimony of the model indicated by AIC.

**Table A.3.** Model comparisons across three nested models with the same fixed factors (expectation, memory type, time and their interactions) but increased complexity in the included mixed effects for the analysis of pupil responses

| Model (using *lme4* call formatting in *R*) | $R^2_{marginal}$ | $R^2_{conditional}$ | AIC | ICC | Random Effect LRT |
|---|---|---|---|---|---|
| 1) pupil ~ 1 + memory type + expectation + time + memory type:expectation + memory type:time + expectation:time + memory type:expectation:time+( 1 \| subjects) | 0.24 | 0.38 | -3383 | 0.19 | $\chi^2(1) = 61.0$, $p < 0.001$ |
| 2) pupil ~ 1 + memory type + expectation + time + memory type:expectation + memory type:time + expectation:time + memory type:expectation:time+( 1 + memory type \| subjects) | | 0.42 | -3431 | 0.21 | $\chi^2(5) = 57.1$, $p < 0.001$ |
| 3) pupil ~ 1 + memory type + expectation + time + memory type:expectation + memory type:time + expectation:time + memory type:expectation:time+( 1 + memory type + expectation \| subjects) | | 0.46 | -3477* | 0.22 | $\chi^2(4) = 53.7$, $p < 0.001$ |

Note: $R^2_{marginal}$ is the same across all models as the fixed effects remain the same; AIC = Akaike information criterion (smaller numbers indicate preferred model); ICC = intraclass correlation coefficient; *best model fit based on variance explained, significance of improvement after including the random effect and the parsimony of the model indicated by AIC.
